# *In vivo* CRISPR-Cas gene editing with no detectable genome-wide off-target mutations

**DOI:** 10.1101/272724

**Authors:** Pinar Akcakaya, Maggie L. Bobbin, Jimmy A. Guo, Jose M. Lopez, M. Kendell Clement, Sara P. Garcia, Mick D. Fellows, Michelle J. Porritt, Mike A. Firth, Alba Carreras, Tania Baccega, Frank Seeliger, Mikael Bjursell, Shengdar Q. Tsai, Nhu T. Nguyen, Roberto Nitsch, Lorenz M. Mayr, Luca Pinello, Mohammad Bohlooly-Y, Martin J. Aryee, Marcello Maresca, J. Keith Joung

## Abstract

CRISPR-Cas genome-editing nucleases hold substantial promise for human therapeutics^1–5^ but identifying unwanted off-target mutations remains an important requirement for clinical translation^6, 7^. For *ex vivo* therapeutic applications, previously published cell-based genome-wide methods provide potentially useful strategies to identify and quantify these off-target mutation sites^8–12^. However, a well-validated method that can reliably identify off-targets *in vivo* has not been described to date, leaving the question of whether and how frequently these types of mutations occur. Here we describe Verification of *In Vivo* Off-targets **(VIVO)**, a highly sensitive, unbiased, and generalizable strategy that we show can robustly identify genome-wide CRISPR-Cas nuclease off-target effects *in vivo*. To our knowledge, these studies provide the first demonstration that CRISPR-Cas nucleases can induce substantial off-target mutations *in vivo*, a result we obtained using a deliberately promiscuous guide RNA **(gRNA)**. More importantly, we used VIVO to show that appropriately designed gRNAs can direct efficient *in vivo* editing without inducing detectable off-target mutations. Our findings provide strong support for and should encourage further development of *in vivo* genome editing therapeutic strategies.

---

We envisioned VIVO as a two-step strategy to identify CRISPR-Cas nuclease off-target mutations *in vivo* (**Fig. 1a**). In an initial *in vitro* “discovery” step, potential off-target cleavage sites of a nuclease of interest are identified on purified genomic DNA using the recently described CIRCLE-seq method^13^. We used CIRCLE-seq because it is highly sensitive and avoids potential confounding effects associated with cell-based assays performed using surrogate cells in culture^13^. In a second *in vivo* “confirmation” step, off-target sites identified *in vitro* by CIRCLE-seq are examined for evidence of indel mutations in the genomic DNA of target tissues that have been treated with the nuclease. We hypothesized that this two-step strategy might be effective because we have previously shown that the sensitivity of CIRCLE-seq enables it to identify a superset of off-target cleavage sites that includes a subset of loci actually mutagenized in nuclease-treated cells in culture^13^.

**Figure 1.**
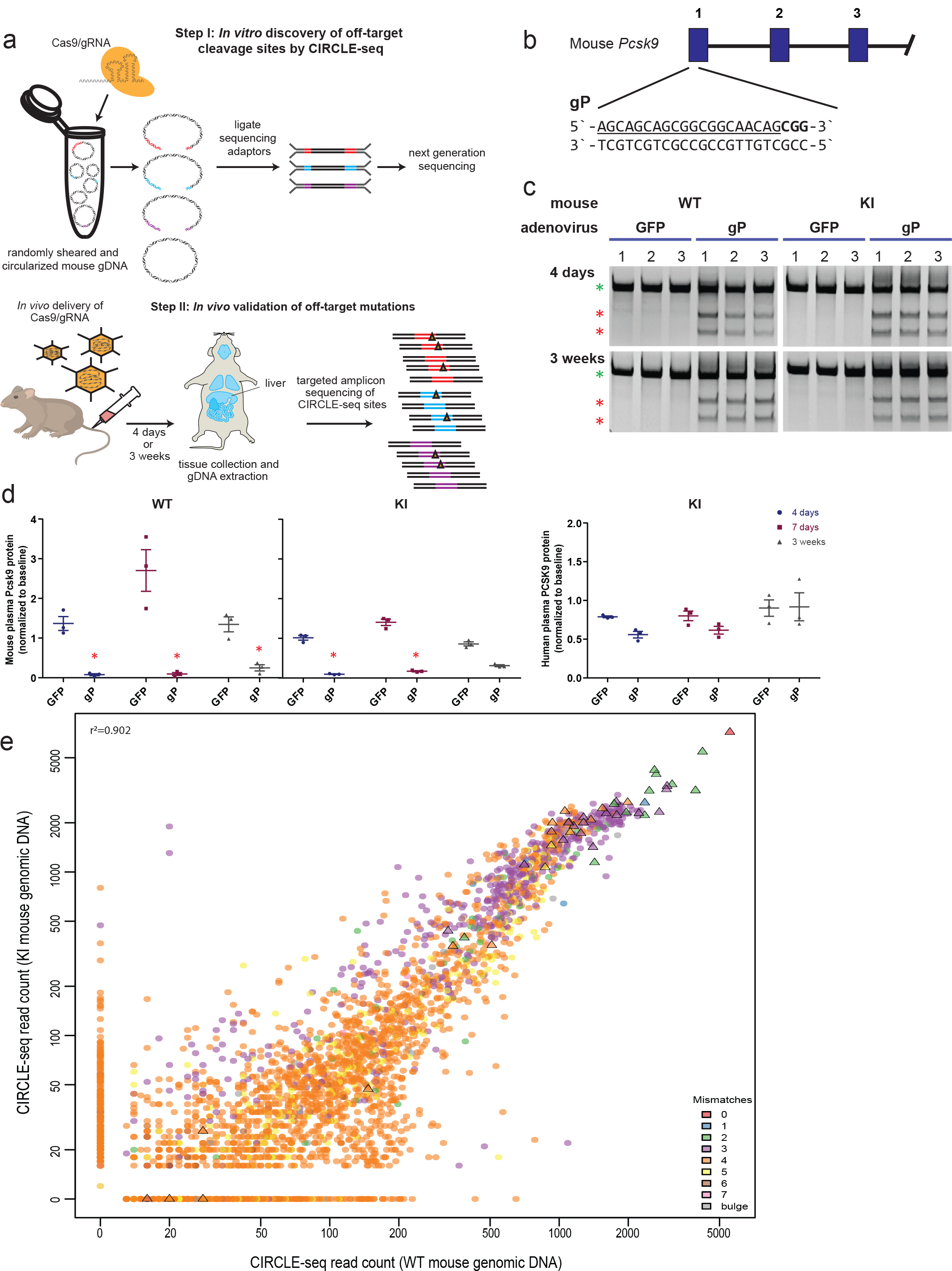
Overview and validation of the Verification of *In Vivo* Off-Targets (VIVO) method. **(a)** Schematic overview of the two-step VIVO method. In Step 1, CIRCLE-seq identifies off-target sites cleaved *in vitro*. In Step 2, sites identified in Step 1 are assessed for indel mutations by targeted amplicon sequencing of sites *in vivo* from the livers of nuclease-treated mice. **(b)** Sequence and location of the SpCas9 promiscuous gP gRNA target site in the mouse *Pcsk9* locus. Blue bars indicate exons. The PAM sequence is in bold and the spacer sequence is underlined. **(c)** Surveyor assay demonstrating efficient *in vivo* modification of the on-target site in mouse liver by the gP gRNA and SpCas9 nuclease. Assays were performed 4 days and 3 weeks after administration of adenoviral vectors encoding gP and SpCas9 (“gP”) or GFP and SpCas9 (“GFP”) using genomic DNA isolated from livers of wild-type C57BL/6N (WT) mice or a derivative strain harbouring a single copy of the human *PCSK9* cDNA gene knocked into the *Rosa26* locus (KI). Green asterisks indicate uncleaved PCR products and red asterisks indicate cleaved PCR products expected following treatment with Surveyor nuclease. **(d)** Plasma mouse Pcsk9 protein levels measured in WT and KI mice and plasma human PCSK9 protein levels measured in KI mice following nuclease treatment. Protein levels were assessed 4 days, 7 days, and 3 weeks following administration of gP or control GFP adenoviral vectors and normalized to baseline levels. Differences between experimental and control groups were determined using two-way ANOVA and Dunnet`s multiple comparisons test with asterisks indicating differences with *p*<0.05. All values are presented as group means, error bars represent standard error of the mean (SEM). **(e)** Scatterplot of CIRCLE-seq read counts for sites identified with gP/SpCas9 on genomic DNA from WT and KI mice. Read counts are shown on a log scale and colors indicate the number of mismatches in each off-target site relative to the on-target site. Sites shown as triangles were chosen for targeted amplicon sequencing.

To test the VIVO strategy, we first designed a *Streptococcus pyogenes* Cas9 (**SpCas9**) gRNA intentionally selected for its high likelihood of inducing multiple off-target mutations in the mouse genome. This promiscuous gRNA (**gP**) targets a sequence within the coding sequence of exon 1 in the mouse *Pcsk9* gene (**Fig. 1b**) and was chosen because it has many closely related sites (i.e., sequences having one, two or three mismatches relative to the on-target site) in the mouse genome (**Online Methods; Extended Data Table 1**). To deliver nucleases efficiently to mouse livers *in vivo* (**Fig. 1a**), we infected cohorts of mice with adenoviral vectors encoding both SpCas9 and gP or with a control virus encoding SpCas9 and GFP. We infected two related strains of mice: a wild-type C57BL/6N strain (hereafter referred to as “**WT**” mice) and a derivative of the C57BL/6N strain that harbors a single copy of a human *PCSK9* cDNA knocked into the *Rosa26* locus (hereafter referred to as “**KI**” mice; Carreras A & Pane SL et al., manuscript in preparation). We observed efficient targeted modification of the on-target *Pcsk9* site (as judged by the Surveyor assay) in liver tissue of both WT and KI mice infected with gP/SpCas9 adenoviral vector at both four days and three weeks post-infection (**Fig. 1c**). Consistent with this, plasma levels of mouse Pcsk9 protein were reduced in both mouse strains treated with the gP/SpCas9 vector compared to those treated with negative control vector at four days, one week, and three weeks post-infection (**Fig. 1d**). Because bio-distribution studies of our adenoviral vectors in mice showed selective delivery to liver (**Extended Data Fig. 1**), we focused our analysis of nuclease-induced indel mutations on this organ.

Having established the efficacy of gP/SpCas9 for *in vivo* on-target *Pcsk9* modification, we conducted the first screening step of VIVO by performing CIRCLE-seq with this nuclease on genomic DNA isolated from the livers of WT and KI mice (**Fig. 1a; Online Methods**). As expected, these experiments identified a large number of *in vitro* off-target cleavage sites for the gP gRNA: 3107 and 2663 sites with the WT and KI mice genomes, respectively (**Fig. 1e** and **Supplementary Table 1**), although these sites represent only a small percentage of all sites in the genome that have seven or fewer mismatches relative to the on-target site (**Extended Data Fig. 2**). As expected, 2368 of the sites were identified in both mouse genomes and showed strong concordance (r^2^ = 0.902) in their numbers of CIRCLE-seq read counts between the two mice strains (we have previously shown that CIRCLE-seq counts provide a semi-quantitative reflection of cleavage efficiency^13^) (**Fig. 1e**). Sites identified in only one genome or the other generally had among the lowest CIRCLE-seq read counts (**Fig. 1e**), consistent with the idea that differential detection might be related to assay limit of detection (although SNPs might also play a role for a subset of sites as well (**Extended Data Table 2**)). The 20 off-target sites with the highest CIRCLE-seq read counts all had three or fewer mismatches in the spacer sequence relative to the on-target site (**Fig. 1e and Supplementary Table 1**). Several off-target sites contained mismatches in the protospacer adjacent motif (**PAM**) sequence, with NAG PAM as the most prevalent (**Fig. 1e** and **Supplementary Table 1**), consistent with data from previously published studies^8, 13–15^.

To perform the second validation step of VIVO, we examined whether gP off-target cleavage sites identified by CIRCLE-seq showed *in vivo* evidence of indel mutations in the livers of WT and KI mouse treated with gP/SpCas9. Because of the very large number of potential gP CIRCLE-seq off-target sites, we performed targeted amplicon sequencing on the following subset of sites on three different mice at both the four day and three week time points: the *Pcsk9* on-target site, 11 “Class I” off-target cleavage sites with the highest CIRCLE-seq read counts (harboring one to three mismatches relative to the on-target site), 17 “Class II” sites with moderate CIRCLE-seq read counts (harboring two to four mismatches), and 17 “Class III” sites with lower CIRCLE-seq read counts (harboring one to six mismatches) (**Figs. 1e** and **2**). Consistent with the Surveyor assay results, we found that the *Pcsk9* on-target site was efficiently mutagenized and remained stable over time in both WT and KI mice, with mean indel frequencies ranging from ^~^23 to 30% (**Fig. 2** and **Supplementary Table 2**). 19 of the 45 potential off-target sites we examined showed significant evidence of indel mutations (mean frequencies ranging from 41.9% to 0.13%) in the livers of both WT and KI mice at four days and three weeks post-infection relative to untreated controls (**Fig. 2** and **Supplementary Table 2**). Notably, higher CIRCLE-seq read count generally correlated well with the probability of finding indel mutations *in vivo* – 11 out of the 11 Class I off-target sites, 5 out of the 17 Class II sites, and 3 out of the 17 Class III sites were shown to harbor indels in mouse liver DNA (**Fig. 2**). In addition, all 19 of these verified *in vivo* off-target sites had three or fewer mismatches relative to the on-target site, with the majority located within gene coding sequences (**Fig. 2**). Three additional sites, including two sites each containing three mismatches in the spacer and one mismatch in the PAM, showed significant indel mutations at only the four-day time point, but were no longer significant by three weeks; the mutation frequencies observed at these sites were around 0.13%, which is close to the limit of detection (0.1%) for this assay. Taken together, our data shows that SpCas9 with a promiscuous gRNA can generate off-target mutations *in vivo* (in some cases with very high frequencies) that are stable over time and that our VIVO approach can also identify even low frequency mutations (as low as 0.13%).

**Figure 2.**
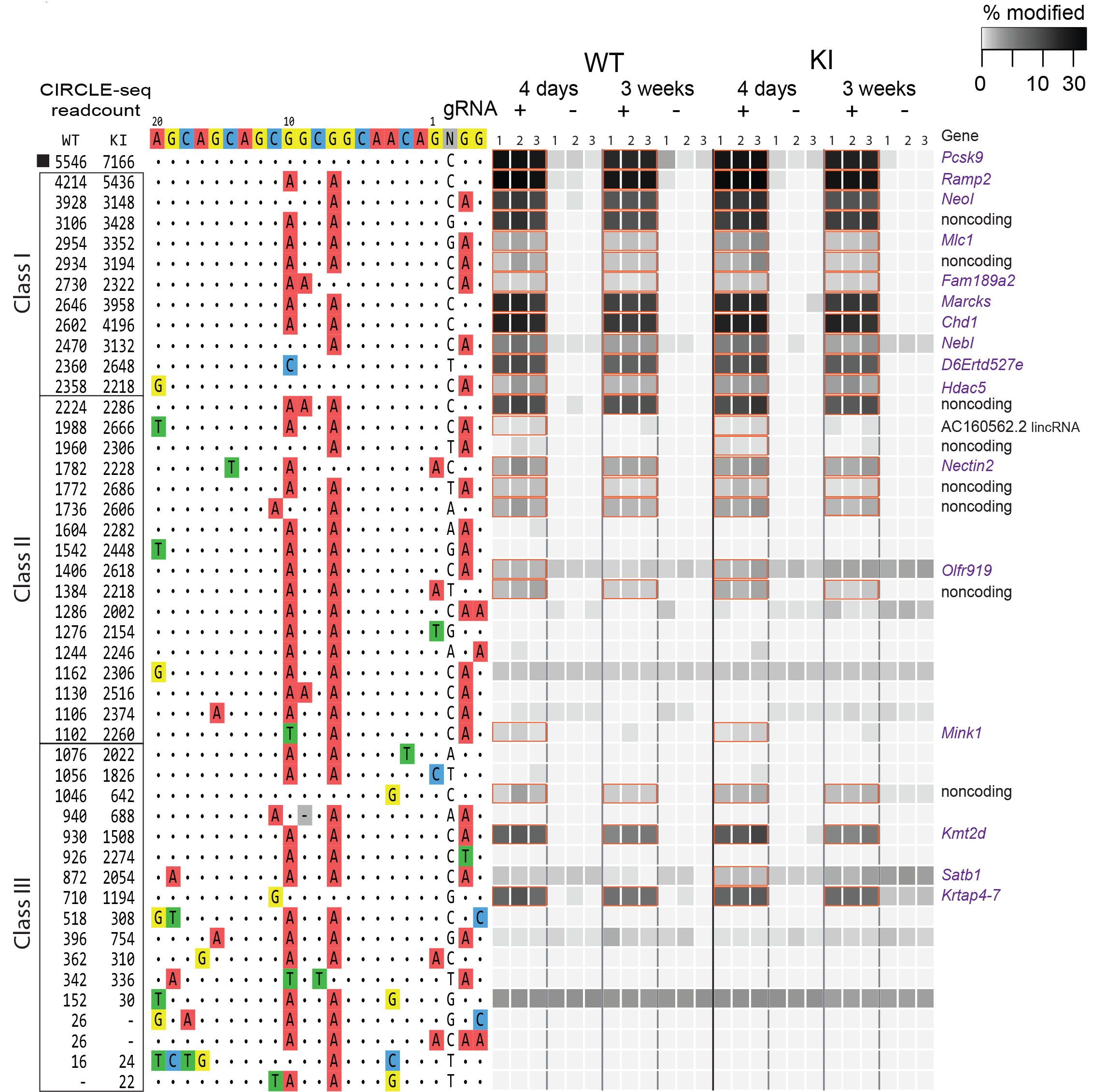
Assessment of *in vivo* off-target indel mutations induced by gP/SpCas9. Indel mutation frequencies determined by targeted amplicon sequencing (using high-throughput sequencing) are presented as heat maps for the gP/SpCas9 on-target site (black square) and Class I, Class II, and Class III off-target sites (defined in the text) identified from CIRCLE-seq experiments. Each locus was assayed in three different mice (1, 2, 3) using genomic DNA isolated from the liver of WT and KI mice treated with experimental adenoviral vector encoding gP/SpCas9 (gRNA +) or control adenoviral vector GFP/SpCas9 (gRNA −). For each site, mismatches relative to the on-target site are shown with colored boxes and bases in the spacer sequence are numbered from 1 (most PAM-proximal) to 20 (most PAM-distal). The number of read counts found for each site from the CIRCLE-seq experiments on WT and KI mouse genomic DNA are shown in the left columns (ranked from highest to lowest based on read counts in the WT genomic DNA CIRCLE-seq experiment). Each box in the heatmap represents a single sequencing experiment. Sites that showed a significant difference between the experimental gRNA + and gRNA−samples are highlighted with an orange outline around the box. Sites identified as having a significant difference and located within the coding sequence of a gene or in non-coding sequence are labeled to the right of the heatmap.

Because actual therapeutic applications would never use a gRNA expected to have a high degree of similarity to many loci in the genome, we sought to characterize the *in vivo* genome-wide off-target profiles of SpCas9 gRNAs designed for sites that are more orthogonal to the mouse genome. Thus, we constructed two additional gRNAs, which we named gM and gMH, to target sites in the mouse *Pcsk9* gene coding sequence (**Fig. 3a**). In contrast to gP, these gRNAs have relatively few closely matched sites (i.e., those with one, two or three mismatches relative to the on-target site) in the C57BL6/N mouse genome (**Extended Data Table 1**). gM has been previously described and characterized in an earlier study^16^. gMH is also predicted to target a site in the human *PCSK9* transgene present in the KI mouse that has one mismatch relative to the on-target site (**Fig. 3a**). To test gM and gMH with SpCas9 *in vivo*, we delivered each gRNA/nuclease pair (and the GFP/SpCas9 negative control) using adenoviral vectors to WT and KI mice. Surveyor assays performed with liver genomic DNA showed efficient and stable modification of the on-target mouse *Pcsk9* site with gM and gMH in WT and KI mice and of the human *PCSK9* transgene site with gMH in the KI mouse (**Fig. 3b**). In addition, we observed significant and stable reductions of mouse PCSK9 protein over time in plasma with gM and gMH in both mouse strains and of human PCSK9 protein in plasma with gMH in the KI mouse (**Fig. 3c**).

**Figure 3.**
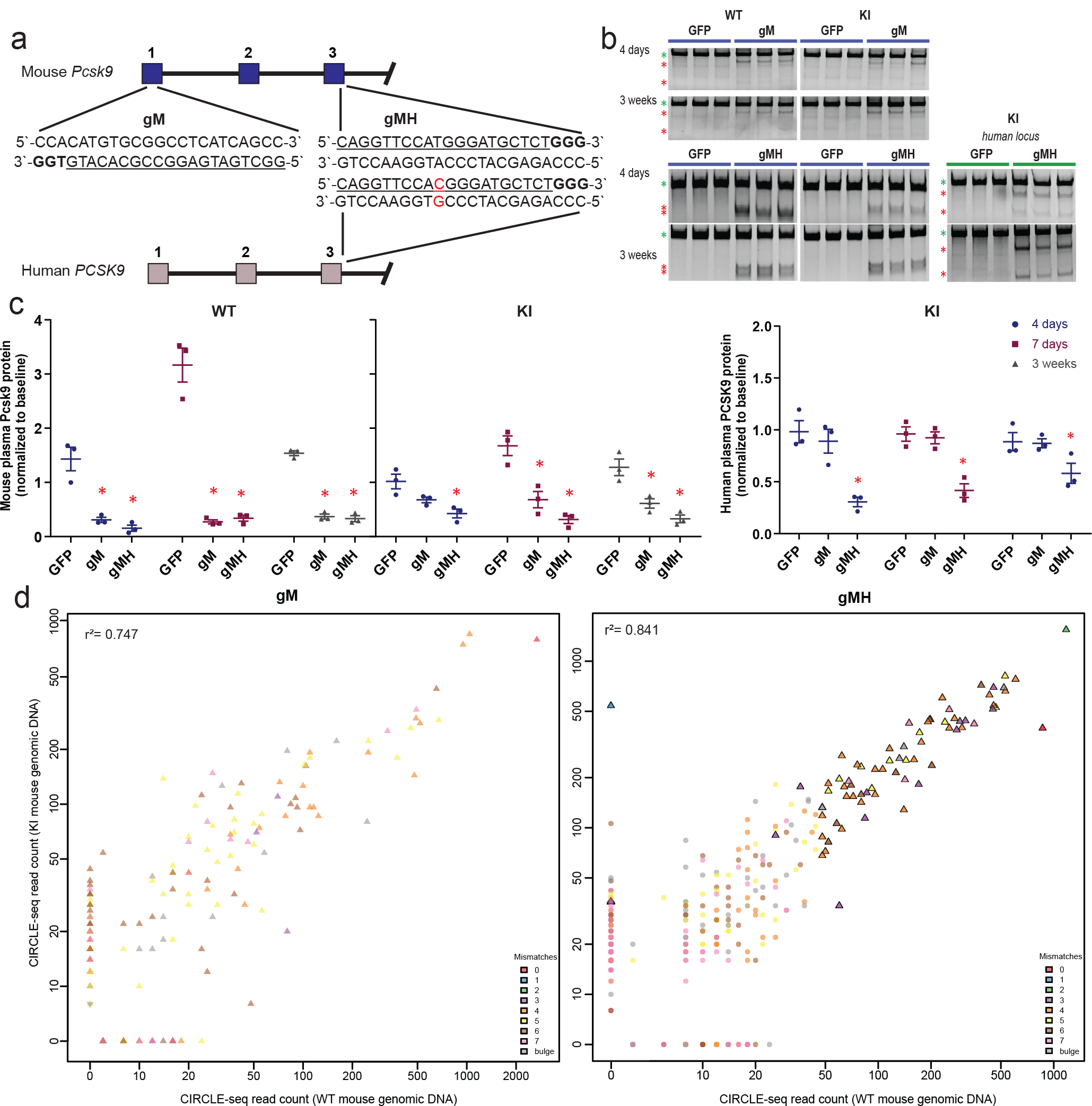
Characterization of *Pcsk9*-targeted gRNAs designed to be orthogonal to the mouse genome. **(a)** Sequence and location of the SpCas9 gM (mouse) and gMH (mouse & human) gRNA target sites in the endogenous mouse *Pcsk9* gene and human *PCSK9* transgene inserted at the mouse *Rosa26* locus. Blue bars indicate exons. PAM sequence for the sites is in bold and the spacer sequence is underlined. The single base position mismatched between the gMH target site and the site in the human *PCSK9* transgene is highlighted in red. **(b)** Surveyor assay demonstrating efficient *in vivo* modification of the on-target endogenous mouse *Pcsk9* site and human PCSK9 transgene in mouse liver. Assays were performed 4 days and 3 weeks following administration of adenoviral vectors encoding gM and SpCas9 (“gP”), gMH and SpCas9 (“gMH”) or GFP and SpCas9 (“GFP”) using genomic DNA isolated from livers of WT and KI mice. Green asterisks indicate uncleaved PCR products and red asterisks indicate cleaved PCR products expected following treatment with Surveyor nuclease. **(c)** Plasma mouse Pcsk9 protein levels measured in WT and KI mice and plasma human PCSK9 protein levels measured in KI mice following CRISPR-Cas nuclease treatment. Plasma protein levels were assessed 4 days, 7 days, and 3 weeks following administration of gP or control GFP adenoviral vectors and normalized to baseline levels at each timepoint. Differences between groups were determined using two-way ANOVA and Dunnet`s multiple comparisons test with asterisks indicating differences with *p*<0.05. All values are presented as group means, error bars represent standard error of the mean (SEM). **(d)** Scatterplot of CIRCLE-seq read counts for sites identified with gM/SpCas9 and gMH/Cas9 on genomic DNA from WT and KI mice. Read counts are shown on a log scale and colors indicate the number of mismatches in each off-target site relative to the on-target site. Sites shown as triangles were chosen for targeted amplicon sequencing.

To carry out the first *in vitro* step of VIVO with gM/SpCas9 and gMH/SpCas9, we performed CIRCLE-seq experiments on genomic DNA from WT and KI mice. We identified a more moderate number of off-target sites than what was obtained with gP/SpCas9: 129 sites for gM on WT mouse DNA, 145 sites for gM on KI mouse DNA, 333 sites for gMH on WT mouse DNA, and 394 sites for gMH on KI mouse DNA (**Fig. 3d, Supplementary Tables 3 and 4**). As we found with the gP gRNA, most of the off-target cleavage sites for gM and gMH were identified in both mouse genomes with good concordance in their CIRCLE-seq read counts (r^2^ = 0.747 and 0.841, respectively) (**Fig. 3d**). Sites found in only one mouse genome or the other generally had low read counts (**Fig. 3d**). As expected, CIRCLE-seq identified the on-target mouse *Pcsk9* in all four experiments and the human *PCSK9* transgene site only in the experiment with gMH on the KI mouse DNA (**Fig. 3d**). Most of the potential off-target sites identified with both gRNAs had three or more mismatches relative to the on-target site, consistent with the higher degree of on-target site orthogonality to the mouse genome.

To conduct the second *in vivo* step of VIVO, we performed targeted amplicon sequencing of potential gM and gMH off-target sites found by CIRCLE-seq using genomic DNA isolated from the livers of adenovirus-treated WT and KI mice. For gM, we comprehensively examined 181 of the 182 off-target CIRCLE-seq cleavage sites (one site could be amplified but not sequenced, see **Online Methods**) and the on-target site. Each site was examined in three WT and three KI
mice, from liver DNA harvested at 4 days or 3 weeks following adenovirus infection. Strikingly, the only site that showed significant indel mutations (relative to mice treated with the control GFP/SpCas9 virus) was the intended gM on-target site (indel frequencies ranging from 12.6% to 18.5%) (**Fig. 4** and **Supplementary Table 5**); no significant off-target indels were identified in any mice at either the four-day or three-week timepoints. For gMH, because CIRCLE-seq identified a large number of potential off-target sites (529 in total in WT and/or KI mouse genomic DNA), we examined the on-target site and a subset of 69 potential off-target sites with up to six mismatches that had the highest CIRCLE-seq read counts. These 69 sites encompassed all but one of the CIRCLE-seq sites that had up to three mismatches (one site could be amplified but not sequenced, see **Online Methods**) and also included the human *PCSK9* transgene site (which bears one mismatch) (**Fig. 3d**). Our choice to test this subset of sites for the gMH gRNA was guided by our findings that none of the 19 *bona fide in vivo* off-target sites we found for gP/SpCas9 had four or more mismatches relative to the on-target site (**Fig. 2**). Among the sites we assessed, we found significant indel mutations at only two sites: the on-target mouse gMH site (indel frequencies ranging from 27.4% to 43.6%) and the human PCSK9 transgene site bearing one mismatch (indel frequencies ranging from 20.4% to 21.7%) (**Extended Data Fig. 3; Supplementary Table 6**).

**Figure 4.**
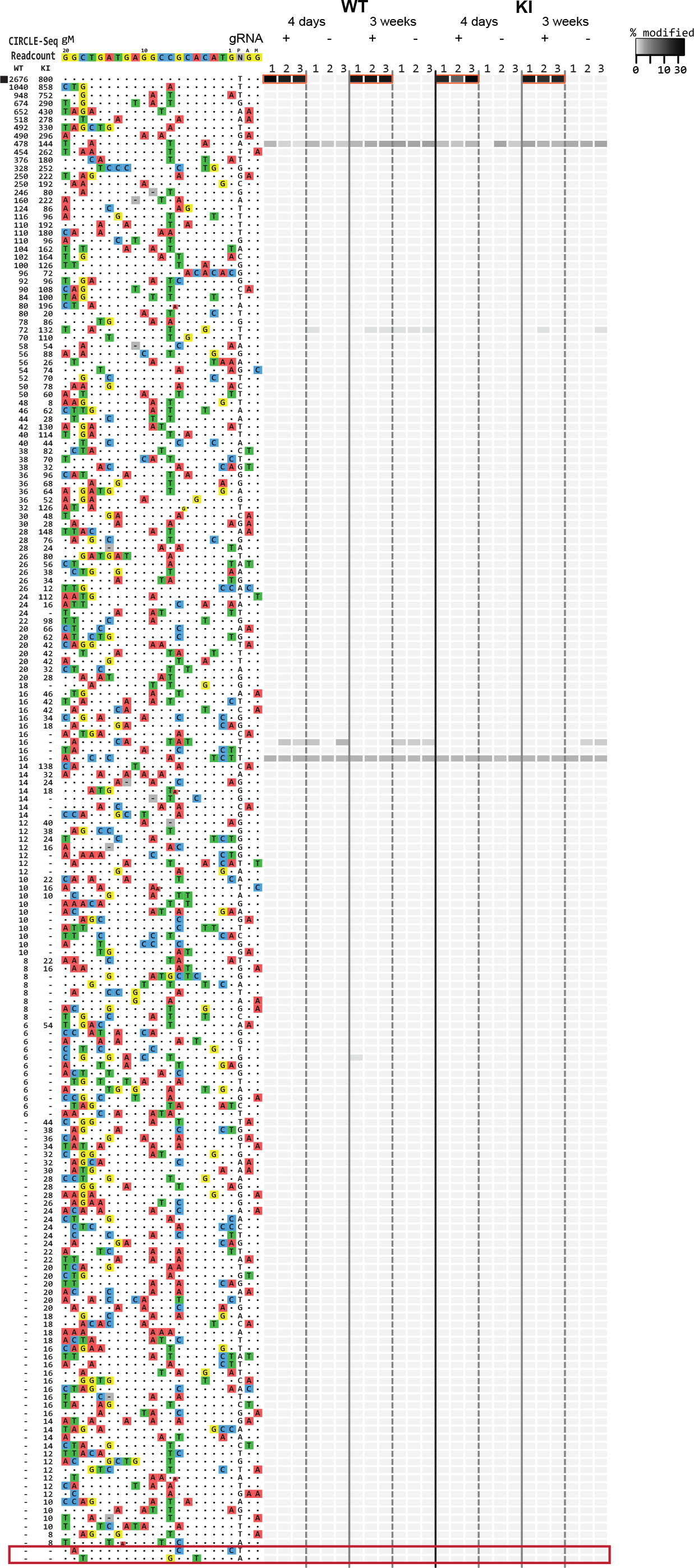
Assessment of *in vivo* off-target indel mutations induced by gM/SpCas9. Indel mutation frequencies determined by targeted amplicon sequencing (using high-throughput sequencing) are presented as heat maps for the gM/SpCas9 on-target site (black square) and 181 off-target sites identified from CIRCLE-seq experiments. Each locus was assayed in three different mice (1, 2, 3) using genomic DNA isolated from the liver of WT and KI mice treated with experimental adenoviral vector encoding gM/SpCas9 (gRNA +) or control adenoviral vector GFP/SpCas9 (gRNA −). For each site, mismatches relative to the on-target site are shown with colored boxes and bases in the spacer sequence are numbered from 1 (most PAM-proximal) to 20 (most PAM-distal). The number of read counts found for each site from the CIRCLE-seq experiments on WT and KI mouse genomic DNA are shown in the left columns (ranked from highest to lowest based on counts in the WT genomic DNA CIRCLE-seq experiment). Each box in the heatmap represents a single sequencing experiment. The single site (the on-target site) that was significantly different between the experimental gRNA + and control gRNA ‒ samples is highlighted with an orange outline around the boxes. Additional closely matched sites in the mouse genome (not identified from the CIRCLE-seq experiments) that were examined for indel mutations are boxed in red at the bottom of the figure.

Given the lack of any observable off-target mutations for gM and gMH *in vivo*, we undertook additional targeted amplicon sequencing to examine the most closely matched sites in the mouse C57BL/6N genome (harboring one, two, or three mismatches in the spacer sequence) that were not identified by our CIRCLE-seq assays (4 sites for gM and 10 sites for gMH) (**Extended Data Table 1**). Three of the gM sites could not be individually selectively amplified and so these sites were assessed together as a pool (Online Methods). Once again, we did not observe significant indel mutations at any of these additional sites in all four sets of treated mice at both time points (**Fig. 4; Extended Data Fig. 3**; and **Supplementary Tables 5** and **6**).

Our results provide, to our knowledge, the first convincing demonstration that CRISPR-Cas nucleases can indeed induce significant off-target mutations *in vivo*. Previous *in vivo* studies have reported no or very few off-target mutations but used approaches that have not been validated to effectively identify these sites *in vivo*^10, 16–31^. Most of these previous reports used computational *in silico* approaches that are known to miss many *bona fide* off-target sites in cell-based systems, making their efficacy questionable in an *in vivo* setting. Three of these studies^17–19^ used the cell-based GUIDE-seq method^8^ to determine what sites to examine *in vivo* and only one found a single low-level off-target (frequency ^~^1%); however, GUIDE-seq was performed on surrogate cells in culture in these experiments, a strategy that may miss off-target sites that occur in the actual target tissue *in vivo*. Our development and validation of VIVO now enables the robust identification of off-target sites *in vivo* (with frequencies as low as ^~^0.1%), with its efficacy likely stemming from the high sensitivity of the *in vitro* CIRCLE-seq method. It is important to emphasize that the *in vivo* detection limit of VIVO, like all existing off-target determination methods, is limited by the current error rate of next-generation sequencing, which sets a floor of about 0.1%. Improving the sensitivity of VIVO will therefore require overcoming this limitation or developing alternatives to targeted amplicon sequencing, an especially important goal given the very large number of cells that would be modified with an *in vivo* CRISPR-Cas nuclease therapeutic.

Our results and the VIVO method define a pathway for assessing and optimizing the *in vivo* genome-wide specificities of CRISPR-Cas nucleases. With VIVO, we have convincingly shown that SpCas9 gRNAs can be designed that fail to induce detectable off-target effects *in vivo* while still efficiently and stably modifying their intended on-target site in mouse liver. Based on these successes, we recommend the initial design of gRNAs with the lowest possible number of closely matched sites (i.e., those with three or fewer mismatches relative to the on-target site) in the target genome, something that can be accomplished using existing *in silico* tools^32^. These gRNAs can then be assessed by CIRCLE-seq to identify those that exhibit a reasonable number (e.g., less than 100-200) of potential off-target sites (VIVO Step 1), which can then be comprehensively examined for indel mutations *in vivo* using targeted amplicon sequencing (VIVO Step 2). For even greater certainty, we also suggest the examination of any closely matched sites (3 or fewer mismatches) in the genome that are not identified by CIRCLE-seq as we did for the gM and gMH gRNAs in this study. Any persistent off-target mutations might be reduced to undetectable levels by using high-fidelity CRISPR-Cas nuclease variants^33–35^. In addition, the delivery of RNAs or ribonucleoprotein complexes (rather than DNA by viral vector as we did in this study) might also further reduce off-target mutations^36^.

We believe that the VIVO approach described here sets a new and important standard for defining off-target effects in future *in vivo* studies. The approach is generalizable and can be used irrespective of the method used to deliver the CRISPR-Cas nucleases. We used adenovirus in this study to achieve efficient liver delivery but VIVO could also be used with other viral or non-viral delivery strategies (e.g., retroviral and lentiviral vectors, lipid nanoparticles). With some minor modification of the CIRCLE-seq protocol, VIVO should also work with other types of nucleases (e.g., zinc finger nucleases, meganucleases, transcription activator-like effector nucleases, CRISPR-Cpf1/Cas12a nucleases). We envision that VIVO might also be extended to non-mammalian organisms (e.g., mosquitoes and plants). One can envision that the VIVO approach could eventually be used in a patient-specific fashion, thereby addressing concerns about the impact of individual genomic variation on off-target profiles^37, 38^. Overall, we believe that results and methods described here should strongly motivate the development and advancement of *in vivo* genome editing therapeutics.

## Online Methods

### Guide RNA design

We identified the promiscuous gP gRNA by searching for an on-target sequence within mouse *Pcsk9* (ENSG00000169174) exons one to three that shows a high number of closely matched sites (two or fewer mismatches to the on-target site) in the mouse genome. To identify gMH, we searched for a gRNA that can cleave both mouse *Pcsk9* and human *PCSK9* and that showed a perfect alignment to mouse *Pcsk9* and up to two nucleotides mismatch to human *PCSK9* (ENSMUSG000000254) at least eight nucleotides distal to the PAM. For both gRNA designs we used AstraZeneca proprietary software as an *in silico* tool, which was developed based on Wellcome Trust Sanger Institute’s codebase (WGE: http://www.sanger.ac.uk/htgt/wge/)^39^ with the addition of the NAG PAM motif as well as NGG in the alignments for potential off-targets^14^. GRCm38/mm10 and GRCh38/hg38 genomes were used as reference for alignments. The gM gRNA targeted to mouse *Pcsk9* has been previously described^16^.

### Adenoviral constructs

Adenoviruses that express SpCas9 and gRNAs (Ad-Cas9-gM, Ad-Cas9-gMH, Ad-Cas9-gP) were generated by Vector Biolabs (Malvern, PA, USA). SpCas9 and gRNAs were expressed from chicken β-actin hybrid (CBh) and U6 promoters, respectively, in a replication-deficient adenoviral-serotype 5 (dE1/E3) backbone. A negative control adenovirus (Ad-Cas9-GFP) that expresses Cas9 and GFP from the CBh and CMV promoters, respectively, but no gRNA was also generated.

### Animal studies

All animal experiments were approved by the AstraZeneca internal committee for animal studies as well as the Gothenburg Ethics Committee for Experimental Animals, (license number: 162-2015+) compliant with EU directives on the protection of animals used for scientific purposes.

C57BL/6N mice (Charles River, Sulzfeld, Germany) were individually housed in a temperature (21±2°C) and humidity (55±15%) controlled room with a 12:12 hours light:dark cycle. R3 diet (Lactamin AB, Stockholm, Sweden) and tap water were provided ad *libitum*. Cage bedding and enrichments include spen chips, shredded paper, gnaw sticks and a plastic house.

Humanized hypercholesterolemia mouse model was generated by liver-specific overexpression of human PCSK9 in C57BL/6N mice (Carreras A & Pane SL et al., manuscript in preparation). Briefly, we cloned human PCSK9 cDNA from the HEK293 genome downstream of the mouse albumin promoter isolated from Albumin-Cre mice (B6.Cg-Tg(Alb-cre)21Mgn/J, the Jackson Laboratory, Bar Harbor, MN) in a vector designed to target the mouse *Rosa26* locus. Mouse embryonic stem cells were electroporated with linearized plasmid. Clones that have integration in *Rosa26* locus were selected with neomycin resistance and used for mouse generation. For *in vivo Pcsk9* gene editing, nine- to eleven-week-old male mice received a tail vein injection with a dose of 1 × 109 infection units (IFU) adenovirus (Ad-Cas9-gM, Ad-Cas9-gMH, Ad-Cas9-gP or Ad-Cas9-GFP) in 200 μl diluted with phosphate-buffered saline. Peripheral blood was sampled before virus administration (baseline), a week after virus administration, and at termination (four days or three weeks after virus administration). Animals were euthanized by cardiac puncture under isoflurane anesthesia at the experimental endpoint. The organs including liver, spleen, lungs, kidney, muscle, brain, and testes were dissected, snap-frozen in liquid nitrogen and stored at −80 until further analyses.

Genomic DNA from frozen tissues was isolated using the Gentra Puregene Tissue kit (Qiagen, Hilden, Germany). *In vivo* gene editing efficiency was evaluated using Surveyor mismatch cleavage assay (Integrated DNA Technologies, BVBA, Leuven, Belgium) and targeted deep sequencing (primers are listed in **Supplementary Tables 2, 5, and 6**).

### Assessment of human PCSK9/mouse Pcsk9 protein levels in plasma

Peripheral blood was collected in EDTA-coated capillary tubes from *vena saphena* during the course of the study and by cardiac puncture at the time of termination. Samples were kept on ice for up to 2 hours prior to extraction of plasma by centrifugation at 10,000 rpm for 20 min at 4°C. Plasma was stored at −80°C until the samples were analyzed. Plasma human PCSK9 and mouse Pcsk9 levels were determined with a standard ELISA kit (DPC900 and MPC900; R&D Systems, Minneapolis, MN, USA) according to the manufacturer’s instructions. Prior to the assay, plasma samples were diluted 1:1000 and 1:800 for human PCSK9 and mouse Pcsk9, respectively.

### Reference Genome for CIRCLE-seq, CRISPResso and Cas-OFFinder

We used build 38 of the C57BL/6NJ genome, sequenced by the Sanger Mouse Genomes Project (http://csbio.unc.edu/CCstatus/pseudo2/C57BL6NJ_b38_f.fa.gz). We included the human PCSK9 gene DNA sequence inserted into the mouse genome as an extra chromosome in the reference and named it “chrPCSK9KI”.

### CIRCLE-seq

CIRCLE-seq was performed experimentally as previously described^13^. Data was processed using v1.1 of the CIRCLE-Seq analysis pipeline^40^ (https://github.com/tsailabSJ/circleseq) with parameters: “window_size: 3; mapq_threshold: 50; start_threshold: 1; gap_threshold: 3; mismatch_threshold: 7; merged_analysis: False, variant_analysis: True”.

### Targeted amplicon deep sequencing

Genomic DNA from liver tissue of adenovirus injected mice was extracted at 4 days and 3 weeks post-treatment for indel analysis. As detailed in the text, to validate off-targets identified by CIRCLE-seq for gP we selected sites with read counts above 50% of the on-target and a variety of lower-ranked sites (containing up to 6 mismatches relative to the on-target) for targeted deep sequencing. In addition, to rule out that CIRCLE-seq was not missing potential off-target sites identified by *in silico* tools, we sequenced all sites containing up to 3 mismatches identified by Cas-OFFinder^32^ for gM and gMH. All sites analyzed were amplified from 150 ng of input genomic DNA (approximately 5 × 10^4^ genomes) with Phusion Hot Start Flex DNA polymerase (New England Biolabs). PCR products were purified using magnetic beads made as previously described^41^, quantified using a QuantiFlor dsDNA System kit (Promega), normalized to 10 ng/μL per amplicon, and pooled. Pooled samples were end-repaired and A-tailed using an end prep enzyme mix and reaction buffer from NEBNext Ultra II DNA Library Prep Kit for Illumina, and ligated to Illumina TruSeq adapters using a ligation master mix and ligation enhancer from the same kit. Library-prepped samples were then purified with magnetic beads made as previously described^41^, size-selected using PEG/NaCl SPRI solution (KAPA Biosystems), quantified using droplet digital PCR (BioRad), and loaded onto an Illumina MiSeq for deep sequencing. To analyze amplicon sequencing of potential on- and off-targets, we used CRISPResso software^42^ v1.11 (https://github.com/lucapinello/crispresso) with the following parameters: ‘-q 30 -ignore_substitutions --hide_mutations_outside_window_NHEJ’.

For each of the 45 gP off-target sites we examined, we obtained 10,000 or more sequencing reads in at least two samples for treated and control samples at all time points (**Supplementary Table 2**). One potential gM off-target site (chr15:98037617-98037640) and one potential gMH off-target site (chr15:4878177-4878200) were amplified but could not be successfully sequenced. The problematic gM site was amplified with two different sets of primers and both amplicons failed to sequence. The gMH site is in a highly repetitive area with low complexity, and we were unable to differentiate this site from other sites in the genome so the site was removed from analysis. For all of the gM off-target sites we were able to sequence, we obtained 10,000 or more sequencing reads in at least two samples for treated and control samples at all time points (**Supplementary Table 5**). For all but one of the gMH off-target sites we were able to sequence, we obtained 10,000 or more sequencing reads in at least two samples for treated and control samples at all time points (**Supplementary Table 6**). One of the gMH off-target sites we sequenced (chr17:33501685-33501708) did not reach the 10,000 read threshold for any samples or time points but read counts ranged from 2509 to 9149. For the three sites that were identified *in silico* as being highly similar to the gM on-target site but that were not identified by CIRCLE-seq, we were unable to selectively amplify these sites individually due their sequence similarities: chr14:25878231-25878254, chr14:26018001-26018024 and chr14:26157615-26157638. Therefore, for these three sites, the read counts were pooled into one amplicon that encompasses all locations and that is labelled as “chr14:pooled” in **Supplementary Table 5**.

### Cas-OFFinder

Identification of potential off-targets by Cas-OFFinder^32^ (https://github.com/snugel/cas-offinder) was done using the off-line version allowing up to 7 mismatches and non-canonical PAMs. We then restricted the output to the sites with at most 6 mismatches in the spacer and at most 1 mismatch in the PAM.

### Non-reference genetic variation

samtools mpileup 1.3.1^43^ was used to discover non-reference genetic variation at the off-target sites identified by CIRCLE-seq. Positions with a genotype quality score greater than 5 and depth of at least 3 were considered as potential variants if they did not fall adjacent to the cleavage site or at the edge of the reads and were not located in a highly repetitive region with poor mapping quality.

### Statistical analysis

Data visualization and statistical analyses for plasma protein measurements were performed using GraphPad Prism 7.02. Protein levels after the adenoviral administration were normalized to baseline levels, and values for gRNA treatment groups were compared with the control treatment group. Comparisons between groups were performed using two-way ANOVA and Dunnett`s multiple comparisons test. *p*≤0.05 was considered to be statistically significant.

### Statistical analysis of targeted amplicon deep sequencing data

*p*-values were obtained by fitting a negative binomial generalized linear model (function MASS:glm.nb in R version 3.4.2) to the control and nuclease-treated samples for each evaluated site and adjusted for multiple comparisons using the Benjamini and Hochberg method (function p.adjust in R version 3.4.2). Multiple testing adjustment was performed within strata defined by guide, mouse background, and timepoint. We considered the indel percentage in the gRNA/SpCas9-treated replicates to be significantly greater than the indel percentage in the GFP/SpCas9-treated controls if the adjusted *p*-value was less than 0.1, the model parameter is greater than zero, and the median indel frequency of the treated replicates is greater than 0.1%.

### Data availability

The data sets generated and analyzed as part of this study are available upon request from the corresponding authors and will be available publicly upon publication.

## Acknowledgements

J.K.J. is supported by the Desmond and Ann Heathwood MGH Research Scholar Award. J.K.J., M.L.B., and J.G. were supported by a sponsored research agreement with AstraZeneca. L.P. is supported by a National Human Genome Research Institute (NHGRI) Career Development Award (R00HG008399). J.K.J., M.J.A. and J.M.L. are supported by a National Institutes of Health Maximizing Investigators’ Research Award (MIRA) (R35 GM118158). J.K.J., L.P. and M.K.C. are supported by the Defense Advanced Research Projects Agency (HR0011-17-2-0042). We thank Mike Snowden, Stefan Platz and Steve Rees for resource allocation from AstraZeneca Research Funds. We thank Jonathan Y. Hsu for helpful discussions and input.

## Author Contributions

P.A., M.J.P., A.C., T.B., M.B., M.M., and R.N. performed in vivo experiments. M.L.B, J.A.G., S.Q.T., and N.T.N. performed the CIRCLE-seq and targeted amplicon sequencing off-target experiments. J.M.L., M.K.C., S.P.G., L.P., and M.J.A. performed bioinformatic and computational analysis of the off-target experiments. M.A.F. generated AstraZeneca proprietary software for gRNA identification. P.A., M.L.B., S.Q.T., M.M., M.B-Y., R.N., M.D.F, L.M.M., F.S., and J.K.J. conceived of and designed the study. P.A., M.L.B., M.M., and J.K.J. organized and supervised experiments. P.A., M.L.B., J.A.G., M.M., and J.K.J. prepared the manuscript with input from all authors.

---

**Extended Data Fig. 1.**
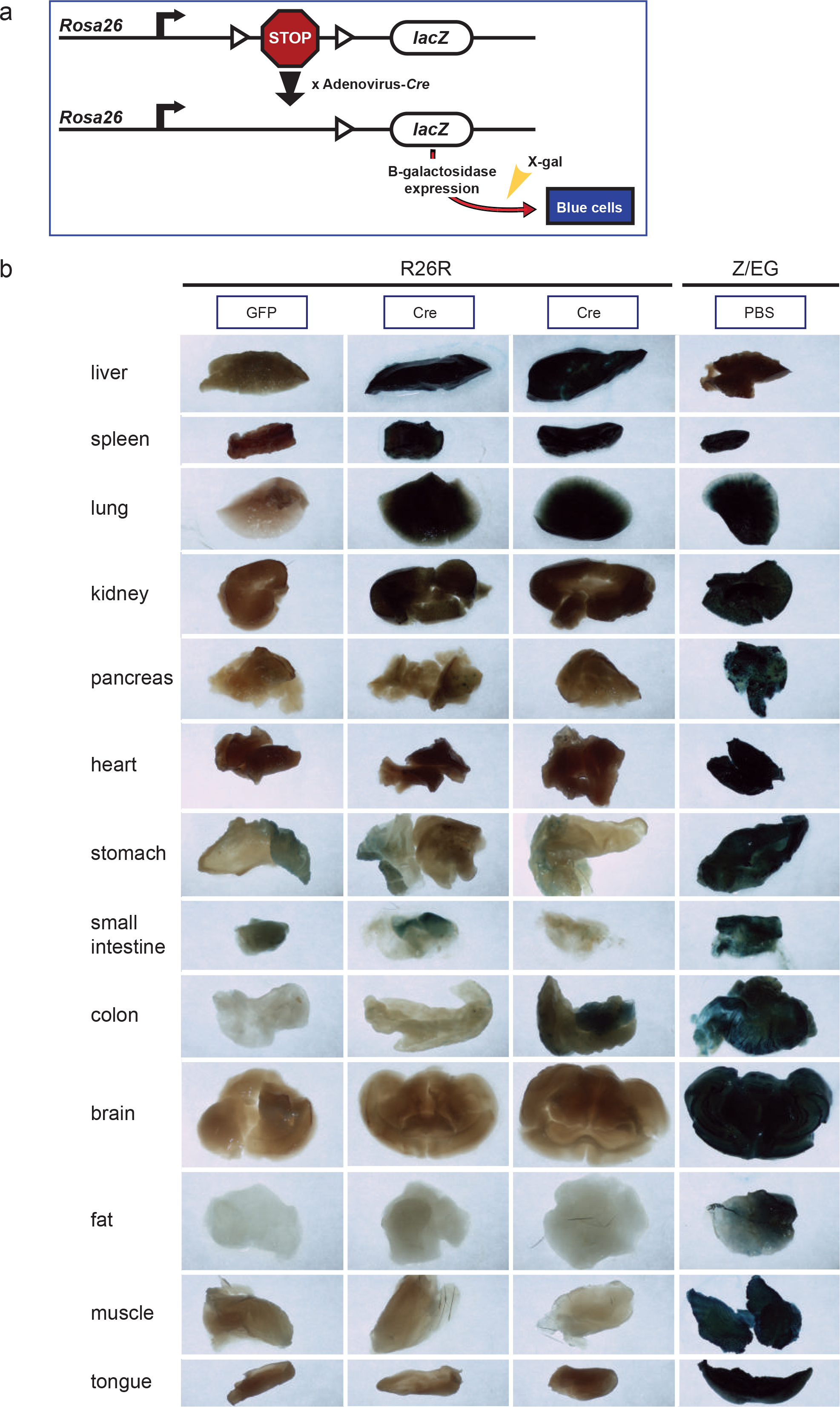
Bio-distribution studies of adenovirus-serotype 5 in mice. (**a**) Schematic of integrated reporter construct in R26R mice used to assess delivery of Cre recombinase using adenovirus-serotype 5 vector. Cre-mediated excision of a *lox*P-flanked transcriptional stop signal upstream of a *lac*Z gene results in expression of beta-galactosidase enzyme. Beta-galactosidase expression can be quantified by staining dissected tissues with X-gal, a compound that turns blue when cleaved by this enzyme. (**b**) Quantification of beta-galactosidase expression in sections of various dissected organs from two R26R mice intravenously injected with adenovirus-serotype 5 vector encoding Cre. Matched organs sections from a R26R mouse intravenously injected with an adenovirus-serotype 5 vector encoding GFP were used to determine background staining levels and serve as a negative control. Matched organ sections from Z/EG mice that constitutively express lacZ (beta-galactosidase) and intravenously injected with PBS (rather than adenovirus) were used to provide positive staining controls. All mice were evaluated one week after adenovirus or PBS injection.

**Extended Data Fig. 2.**
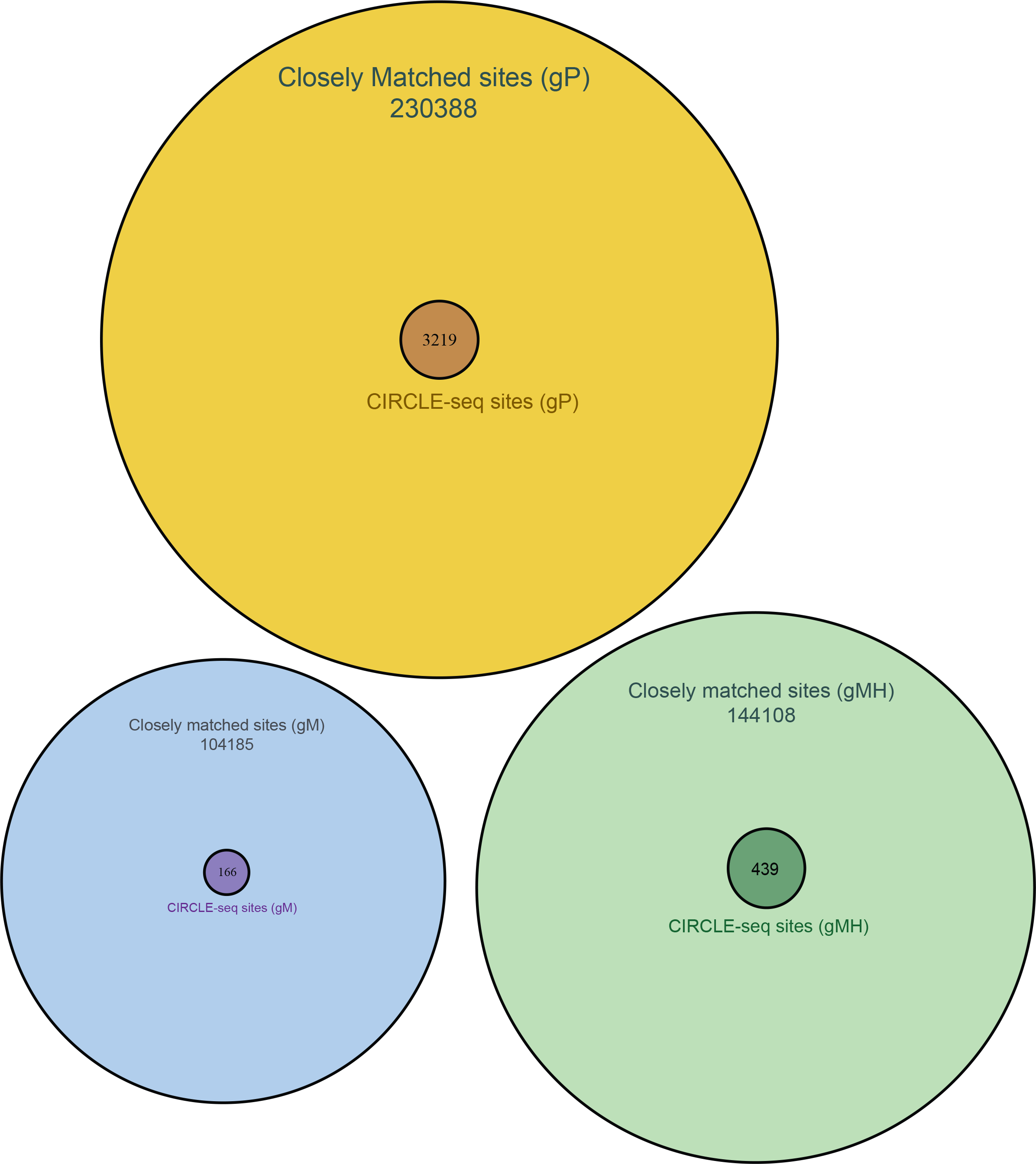
Venn diagrams comparing off-target cleavage sites in mouse genomic DNA identified by CIRCLE-seq experiments with closely matched sites (up to six mismatches relative to the on-target site) in the mouse genome identified *in silico* by Cas-OFFinder. Diagrams are shown for SpCas9 gRNAs gP, gM, and gMH.

**Extended Data Fig. 3.**
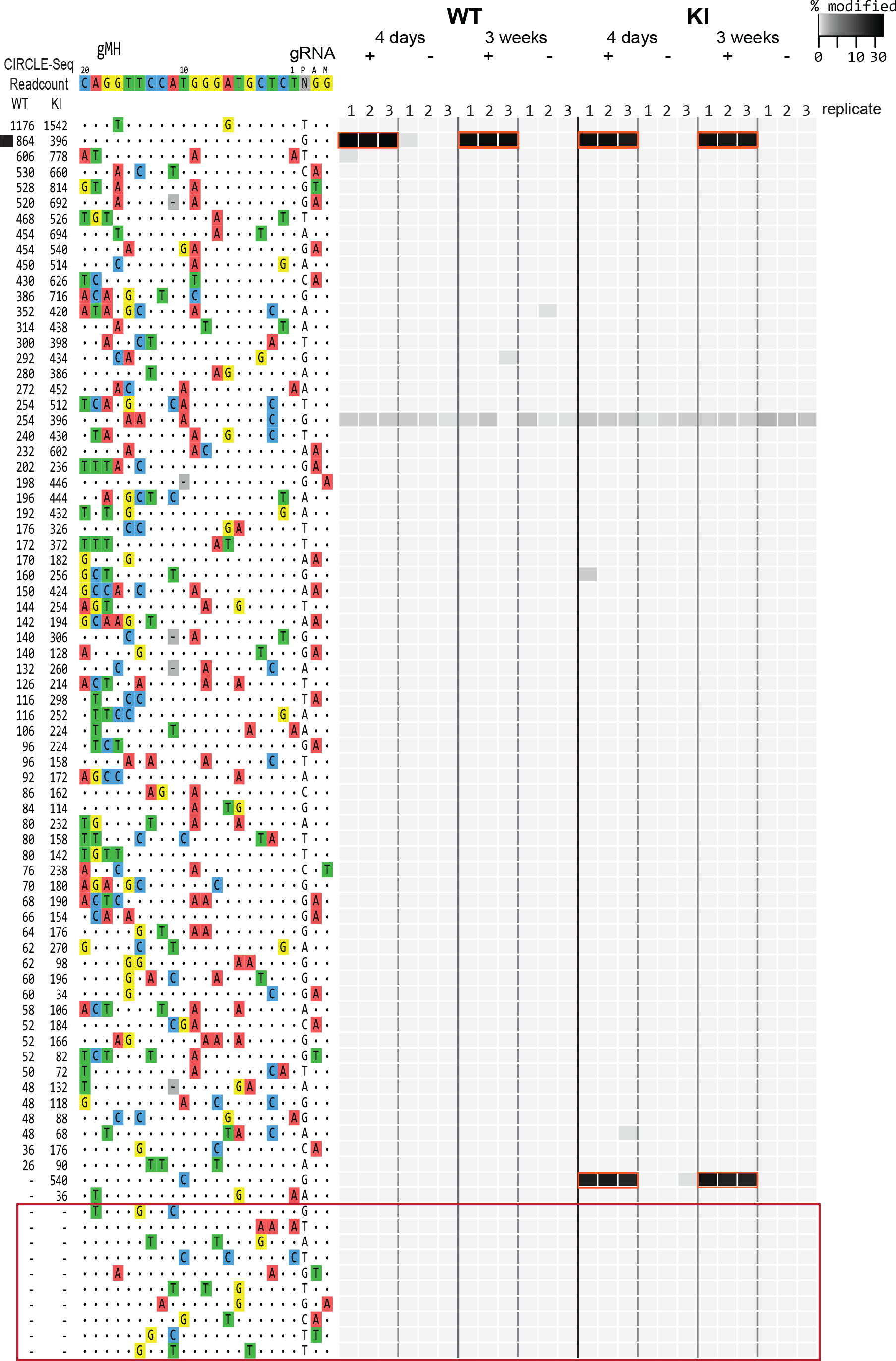
Assessment of *in vivo* off-target indel mutations induced by gMH/SpCas9. Indel mutation frequencies determined by targeted amplicon sequencing (using high-throughput sequencing) are presented as heat maps for the gMH/SpCas9 on-target site (black square) and 63 off-target sites identified from CIRCLE-seq experiments. Each locus was assayed in three different mice (1, 2, 3) using genomic DNA isolated from the liver of WT and KI mice treated with experimental adenoviral vector encoding gM/SpCas9 (gRNA +) or control adenoviral vector GFP/SpCas9 (gRNA −). For each site, mismatches relative to the on-target site are shown with colored boxes and bases in the spacer sequence are numbered from 1 (most PAM-proximal) to 20 (most PAM-distal). The number of read counts found for each site from the CIRCLE-seq experiments on WT and KI mouse genomic DNA are shown in the left columns (ranked from highest to lowest based on counts in the WT genomic DNA CIRCLE-seq experiment). Each box in the heatmap represents a single sequencing experiment. Sites that were significantly different between the experimental gRNA + and control gRNA ‒ samples are highlighted with a red outline around the boxes. Additional closely matched sites in the mouse genome (not identified from the CIRCLE-seq experiments) that were examined for indel mutations are boxed in red at the bottom of the figure.

**Extended Data Table 1:** Numbers of off-target sites for gP, gM, and gMH gRNAs identified by Cas-OFFinder (*in silico*) and CIRCLE-seq (experimental). For sites identified by CIRCLE-seq, the numbers of sites found in the WT and KI mouse are shown and the total number of unique sites found between the two experiments.

|  |  | Sites with canonical NGG PAM |  |  |  |  |  |  |  |
| --- | --- | --- | --- | --- | --- | --- | --- | --- | --- |
|  |  | Number of spacer mismatches |  |  |  |  |  |  |  |
| gRNA | Method | 0 | 1 | 2 | 3 | 4 | 5 | 6 | 7 |
| gP | Cas-OFFinder | 1 | 5 | 41 | 355 | 1073 | 3347 | 21900 | 94051 |
|  | CIRCLE-seq (total WT&KI) | 1 | 5 | 38 | 231 | 121 | 44 | 15 | 4 |
|  | CIRCLE-seq (WT mouse) | 1 | 5 | 38 | 226 | 117 | 43 | 13 | 2 |
|  | CIRCLE-seq (KI mouse) | 1 | 5 | 38 | 218 | 93 | 25 | 11 | 4 |
| gM | Cas-OFFinder | 1 | 0 | 0 | 8 | 77 | 780 | 8315 | 55093 |
|  | CIRCLE-seq (total WT&KI) | 1 | 0 | 0 | 4 | 18 | 36 | 32 | 25 |
|  | CIRCLE-seq (WT mouse) | 1 | 0 | 0 | 4 | 17 | 26 | 23 | 17 |
|  | CIRCLE-seq (KI mouse) | 1 | 0 | 0 | 3 | 17 | 30 | 23 | 14 |
| gMH | Cas-OFFinder | 1 | 1 | 1 | 15 | 178 | 1609 | 10992 | 55363 |
|  | CIRCLE-seq (total WT&KI) | 1 | 1 | 1 | 9 | 52 | 65 | 82 | 159 |
|  | CIRCLE-seq (WT mouse) | 1 | 0 | 1 | 8 | 46 | 46 | 45 | 81 |
|  | CIRCLE-seq (KI mouse) | 1 | 1 | 1 | 9 | 43 | 53 | 64 | 102 |
|  |  | Sites with PAM harboring single mismatch |  |  |  |  |  |  |  |
|  |  | Number of spacer mismatches |  |  |  |  |  |  |  |
| gRNA | Method | 0 | 1 | 2 | 3 | 4 | 5 | 6 |  |
| gP | Cas-OFFinder | 0 | 16 | 370 | 12028 | 9828 | 20097 | 70495 |  |
|  | CIRCLE-seq (total WT&KI) | 0 | 14 | 279 | 2160 | 271 | 31 | 6 |  |
|  | CIRCLE-seq (WT mouse) | 0 | 14 | 272 | 1648 | 164 | 13 | 2 |  |
|  | CIRCLE-seq (KI mouse) | 0 | 14 | 271 | 1904 | 262 | 31 | 4 |  |
| gM | Cas-OFFinder | 0 | 0 | 1 | 20 | 423 | 4911 | 34722 |  |
|  | CIRCLE-seq (total WT&KI) | 0 | 0 | 1 | 3 | 18 | 18 | 10 |  |
|  | CIRCLE-seq (WT mouse) | 0 | 0 | 0 | 3 | 15 | 16 | 8 |  |
|  | CIRCLE-seq (KI mouse) | 0 | 0 | 1 | 3 | 14 | 8 | 4 |  |
| gMH | Cas-OFFinder | 0 | 0 | 7 | 80 | 907 | 8285 | 67109 |  |
|  | CIRCLE-seq (total WT&KI) | 0 | 0 | 3 | 19 | 21 | 9 | 17 |  |
|  | CIRCLE-seq (WT mouse) | 0 | 0 | 3 | 18 | 18 | 7 | 9 |  |
|  | CIRCLE-seq (KI mouse) | 0 | 0 | 3 | 16 | 9 | 5 | 13 |  |

**Extended Data Table 2:**
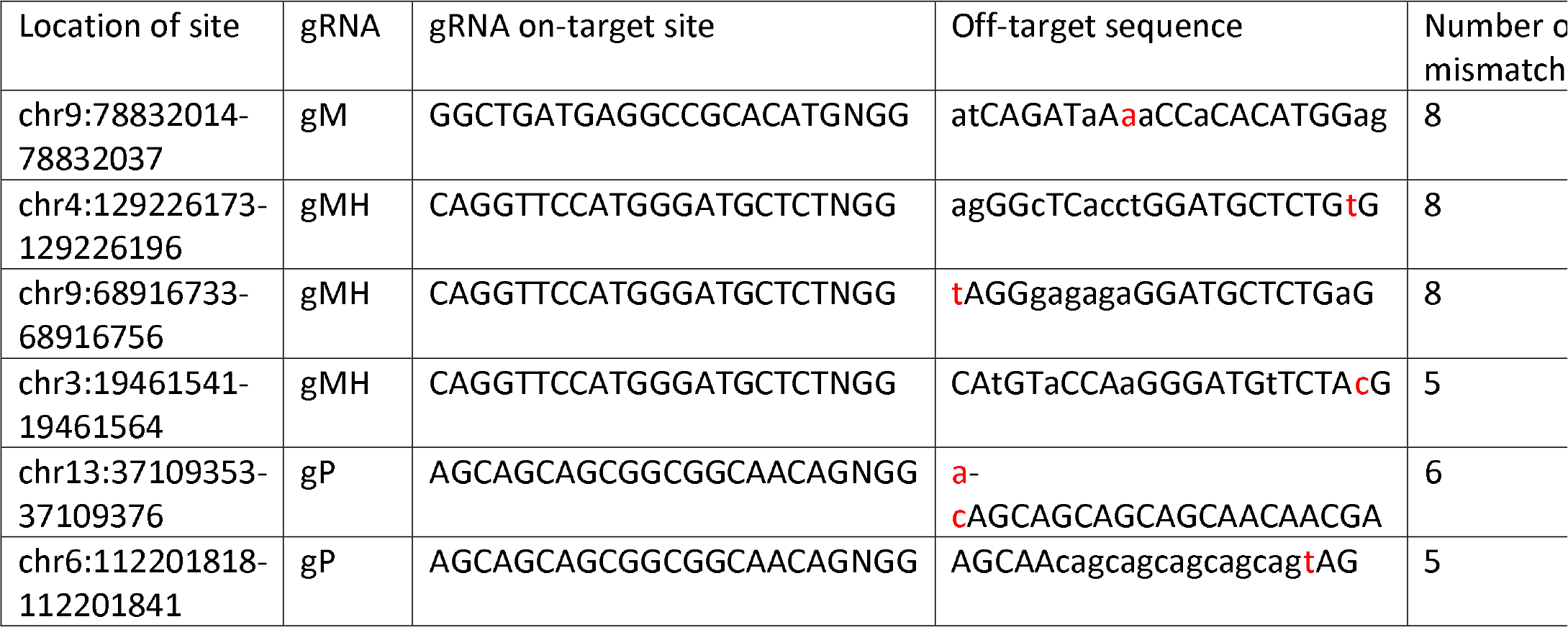
Off-target sites identified by CIRCLE-seq for gP, gM, and gMH that exhibit single nucleotide polymorphisms (SNPs). Single nucleotide mismatches that differ from the C57/BL6N mouse strain are in red.

## Supplementary Table Legends

**Supplementary Table 1.** Off-target cleavage sites identified by CIRCLE-seq for SpCas9 and the gP gRNA. Chromosomal coordinates are provided followed by the CIRCLE-seq readcount for off-target sites found in WT mouse gDNA, the off-target sequence, number of mismatches, bulge mismatch sequence, the score of the number of mismatches with bulges, and the CIRCLE-seq read count for off-targets identified in KI mouse gDNA.

**Supplementary Table 2:** Potential off-target sites for gP/SpCas9 analyzed by targeted amplicon deep sequencing from the livers of WT and KI mice treated with adenoviral vectors. The first column contains the chromosomal site, followed by the MiSeq run and the FASTQ files the amplicon sequencing data was retrieved from. The cell-type, time point, treatment, and replicate follow. The next column gives total read counts obtained for the amplicon followed by the total number of indels found by CRISPResso and the percent of reads containing indels found by CRISPResso at the gDNA cut site. Run refers to the alignment of amplicon to the reference genome and block refers to the run that amplicon was done in. The CIRCLE-seq read counts for each site are also provided and followed by the off-target sequence, mismatch score and sequence of the primers used to amplify each amplicon.

**Supplementary Table 3:** Off-target cleavage sites identified by CIRCLE-seq for SpCas9 and the gM gRNA. Chromosomal coordinates are provided followed by the CIRCLE-seq read count for off-target sites found in WT mouse gDNA, the off-target sequence, number of mismatches, bulge mismatch sequence, the score of the number of mismatches with bulges, and the CIRCLE-seq read count for off-targets identified in KI mouse gDNA.

**Supplementary Table 4:** Off-target cleavage sites identified by CIRCLE-seq for SpCas9 and the gMH gRNA. Chromosomal coordinates are provided followed by the CIRCLE-seq read count for off-target sites found in WT mouse gDNA, the off-target sequence, number of mismatches, bulge mismatch sequence, the score of the number of mismatches with bulges, and the CIRCLE-seq read count for off-targets identified in KI mouse gDNA.

**Supplementary Table 5:** Potential off-target sites for gM/SpCas9 analyzed by targeted amplicon deep sequencing from the livers of WT and KI mice treated with adenoviral vectors. The first column contains the chromosomal site, followed by the MiSeq run and the FASTQ files the amplicon sequencing data was retrieved from. The cell-type, time point, treatment, and replicate follow. The next column gives total read count obtained for the amplicon followed by the total number of indels found by CRISPResso and the percent of reads containing indels found by CRISPResso at the gDNA cut site. Run refers to the alignment of amplicon to the reference genome and block refers to the run that amplicon was done in. The CIRCLE-seq read counts for each site are also provided and followed by the off-target sequence, mismatch score and sequence of the primers used to amplify each amplicon.

**Supplementary Table 6:** Potential off-target sites for gMH/SpCas9 analyzed by targeted amplicon deep sequencing from the livers of WT and KI mice treated with adenoviral vectors. The first column contains the chromosomal site, followed by the MiSeq run and the FASTQ files the amplicon sequencing data was retrieved from. The cell-type, time point, treatment, and replicate follow. The next column gives total read count obtained for the amplicon followed by the total number of indels found by CRISPResso and the percent of reads containing indels found by CRISPResso at the gDNA cut site. Run refers to the alignment of amplicon to the reference genome and block refers to the run that amplicon was done in. The CIRCLE-seq read counts for each site are also provided and followed by the off-target sequence, mismatch score and sequence of the primers used to amplify each amplicon.

## References

1. Musunuru, K. The Hope and Hype of CRISPR-Cas9 Genome Editing: A Review. JAMA cardiology 2, 914–919 (2017).

2. Fellmann, C., Gowen, B.G., Lin, P.C., Doudna, J.A. & Corn, J.E. Cornerstones of CRISPR-Cas in drug discovery and therapy. Nat Rev Drug Discov 16, 89–100 (2017).

3. Komor, A.C., Badran, A.H. & Liu, D.R. CRISPR-Based Technologies for the Manipulation of Eukaryotic Genomes. Cell 168, 20–36 (2017).

4. Koo, T. & Kim, J.S. Therapeutic applications of CRISPR RNA-guided genome editing. Brief Funct Genomics 16, 38–45 (2017).

5. Maeder, M.L. & Gersbach, C.A. Genome-editing Technologies for Gene and Cell Therapy. Mol Ther (2016).

6. Kanchiswamy, C.N., Maffei, M., Malnoy, M., Velasco, R. & Kim, J.S. Fine-Tuning Next-Generation Genome Editing Tools. Trends Biotechnol 34, 562–574 (2016).

7. Tsai, S.Q. & Joung, J.K. Defining and improving the genome-wide specificities of CRISPR-Cas9 nucleases. Nat Rev Genet 17, 300–312 (2016).

8. Tsai, S.Q. et al. GUIDE-seq enables genome-wide profiling of off-target cleavage by CRISPR-Cas nucleases. Nat Biotechnol 33, 187–197 (2015).

9. Frock, R.L. et al. Genome-wide detection of DNA double-stranded breaks induced by engineered nucleases. Nat Biotechnol 33, 179–186 (2015).

10. Ran, F.A. et al. In vivo genome editing using Staphylococcus aureus Cas9. Nature 520, 186–191 (2015).

11. Gao, L. et al. Engineered Cpf1 variants with altered PAM specificities. Nat Biotechnol (2017).

12. Wang, X. et al. Unbiased detection of off-target cleavage by CRISPR-Cas9 and TALENs using integrase-defective lentiviral vectors. Nat Biotechnol 33, 175–178 (2015).

13. Tsai, S.Q. et al. CIRCLE-seq: a highly sensitive in vitro screen for genome-wide CRISPR-Cas9 nuclease off-targets. Nat Methods 14, 607–614 (2017).

14. Hsu, P.D. et al. DNA targeting specificity of RNA-guided Cas9 nucleases. Nat Biotechnol 31, 827–832 (2013).

15. Jiang, W., Bikard, D., Cox, D., Zhang, F. & Marraffini, L.A. RNA-guided editing of bacterial genomes using CRISPR-Cas systems. Nat Biotechnol 31, 233–239 (2013).

16. Ding, Q. et al. Permanent alteration of PCSK9 with in vivo CRISPR-Cas9 genome editing. Circ Res 115, 488–492 (2014).

17. Yin, H. et al. Structure-guided chemical modification of guide RNA enables potent non-viral in vivo genome editing. Nat Biotechnol 35, 1179–1187 (2017).

18. Yin, H. et al. Therapeutic genome editing by combined viral and non-viral delivery of CRISPR system components in vivo. Nat Biotechnol (2016).

19. Gao, X. et al. Treatment of autosomal dominant hearing loss by in vivo delivery of genome editing agents. Nature 553, 217–221 (2018).

20. Bengtsson, N.E. et al. Muscle-specific CRISPR/Cas9 dystrophin gene editing ameliorates pathophysiology in a mouse model for Duchenne muscular dystrophy. Nat Commun 8, 14454 (2017).

21. Long, C. et al. Prevention of muscular dystrophy in mice by CRISPR/Cas9-mediated editing of germline DNA. Science 345, 1184–1188 (2014).

22. Nelson, C.E. et al. In vivo genome editing improves muscle function in a mouse model of Duchenne muscular dystrophy. Science 351, 403–407 (2016).

23. Sanchez-Rivera, F.J. et al. Rapid modelling of cooperating genetic events in cancer through somatic genome editing. Nature 516, 428–431 (2014).

24. Suzuki, K. et al. In vivo genome editing via CRISPR/Cas9 mediated homology-independent targeted integration. Nature 540, 144–149 (2016).

25. Swiech, L. et al. In vivo interrogation of gene function in the mammalian brain using CRISPR-Cas9. Nature biotechnology 33, 102–106 (2015).

26. Wu, Y. et al. Correction of a genetic disease in mouse via use of CRISPR-Cas9. Cell stem cell 13, 659–662 (2013).

27. Tabebordbar, M. et al. In vivo gene editing in dystrophic mouse muscle and muscle stem cells. Science 351, 407–411 (2016).

28. Koo, T. et al. Selective disruption of an oncogenic mutant allele by CRISPR/Cas9 induces efficient tumor regression. Nucleic Acids Res 45, 7897–7908 (2017).

29. Xue, W. et al. CRISPR-mediated direct mutation of cancer genes in the mouse liver. Nature (2014).

30. Han, J. et al. Efficient in vivo deletion of a large imprinted lncRNA by CRISPR/Cas9. RNA Biol 11, 829–835 (2014).

31. Kim, E. et al. In vivo genome editing with a small Cas9 orthologue derived from Campylobacter jejuni. Nat Commun 8, 14500 (2017).

32. Bae, S., Park, J. & Kim, J.S. Cas-OFFinder: a fast and versatile algorithm that searches for potential off-target sites of Cas9 RNA-guided endonucleases. Bioinformatics 30, 1473–1475 (2014).

33. Kleinstiver, B.P. et al. High-fidelity CRISPR-Cas9 nucleases with no detectable genome-wide off-target effects. Nature 529, 490–495 (2016).

34. Slaymaker, I.M. et al. Rationally engineered Cas9 nucleases with improved specificity. Science 351, 84–88 (2016).

35. Chen, J.S. et al. Enhanced proofreading governs CRISPR-Cas9 targeting accuracy. Nature (2017).

36. Zuris, J.A. et al. Cationic lipid-mediated delivery of proteins enables efficient protein-based genome editing in vitro and in vivo. Nat Biotechnol (2014).

37. Lessard, S. et al. Human genetic variation alters CRISPR-Cas9 on-and off-targeting specificity at therapeutically implicated loci. Proc Natl Acad Sci U S A 114, E11257–E11266 (2017).

38. Scott, D.A. & Zhang, F. Implications of human genetic variation in CRISPR-based therapeutic genome editing. Nat Med 23, 1095–1101 (2017).

39. Hodgkins, A. et al. WGE: a CRISPR database for genome engineering. Bioinformatics (Oxford, England) 31, 3078–3080 (2015).

40. Tsai, S.Q., Topkar, V.V., Joung, J.K. & Aryee, M.J. Open-source guideseq software for analysis of GUIDE-seq data. Nat Biotechnol 34, 483 (2016).

41. Rohland, N. & Reich, D. Cost-effective, high-throughput DNA sequencing libraries for multiplexed target capture. Genome Res 22, 939–946 (2012).

42. Pinello, L. et al. Analyzing CRISPR genome-editing experiments with CRISPResso. Nat Biotechnol 34, 695–697 (2016).

43. Li, H. et al. The Sequence Alignment/Map format and SAMtools. Bioinformatics 25, 2078–2079 (2009).

